# DNA damage response signaling is crucial for effective Chikungunya virus replication

**DOI:** 10.1101/2022.04.12.488112

**Authors:** Sanchari Chatterjee, Sameer Kumar, Prabhudutta Mamidi, Ankita Datey, Soumya Sengupta, Chandan Mahish, Eshna Laha, Saikat De, Supriya Suman Keshry, Tapas Kumar Nayak, Soumyajit Ghosh, Sharad Singh, Bharat Bhusan Subudhi, Subhasis Chattopadhyay, Soma Chattopadhyaya

## Abstract

Viruses utilize a plethora of strategies to manipulate the host pathways and hijack its machineries for efficient replication. Several DNA as well as handful of RNA viruses are reported to interact with proteins involved in DNA damage responses (DDR). As the DDR pathways have never been explored in Alphaviruses, this investigation intended to determine the importance of the DDR pathways in CHIKV infection through in vitro, *in vivo* and *ex vivo* models. The study reveals that CHIKV infection activates the Chk2 and Chk1 proteins associated with DDR signaling pathways and increases DNA damage by 95%. Inhibition of both ATM-ATR kinases by ATM/ATR kinase inhibitor (AAKi) shows drastic reduction in viral particle formation in vitro. Next, the treatment of mice with this drug has been shown to reduce the disease score substantially in CHIKV-infected C57BL/6 mice with 71% decrement in the viral copy and the same has been established in hPBMC-derived monocyte-macrophage populations. Additionally, gene silencing of Chk2 and Chk1 reduces viral progeny formation around 73.7% and 78% respectively. Moreover, it has been demonstrated that CHIKV-nsP2 interacts with Chk2 and Chk1 during CHIKV infection and docking analysis depicts the specific amino acids responsible for these interactions. Further, the data suggests that CHIKV infection induces cell cycle arrest in G1 and G2 phases.

In conclusion, this work demonstrated for the first time the mechanistic insight of the induction of DDR pathways by CHIKV that might contribute to the designing of effective therapeutics for the control of this virus infection in future.

**IMPORTANCE:** Viruses being intra-cellular parasite, need several host cell machineries so as to achieve effective replication of their own genome, along with virus-encoded enzymes. One of the strategies is to hijack the DDR pathways. Several DNA as well as handful of RNA viruses interact with the cellular proteins involved in DDR pathways, however, reports with respect to the association of Chk2 and Chk1 in alphavirus infection are scanty. Hence, this study is amongst the first to report that modulation of DDR pathways is crucial for effective CHIKV infection. This work also shows that there is interaction of CHIKV-nsP2 with two crucial host factors, Chk2 and Chk1 for efficient viral infection. Interestingly, CHIKV infection was found to cause DNA damage and arrest cell cycle in G1 and G2 phases to facilitate viral infection. This information might facilitate to develop effective therapeutics for the control of the CHIKV infection in future.

## Introduction

Chikungunya virus (CHIKV) infection is now considered as one of the important public health threat due to the unavailability of vaccines or antiviral treatments. In 1952 CHIKV was discovered in Tanzania, Africa and after that CHIKV endemic have been recorded in several countries of central Africa and southern Asia (1). The recurrence and extravagant spreading of CHIKV in Asia, Europe, and the Americas has considerably escalated research on CHIKV (2–4). The symptoms are mainly high fever, nausea, back pain, rashes over the skin, myalgia, and polyarthralgia. However, it can have a serious imprint on the well being of the affected person for weeks, months, or even years. It belongs to *the* Alphavirus genus *of Togaviridae* family and grouped as Old World alphaviruses. The mode of transmission is by mosquito vectors namely *Aedes agypti* and *Aedes albopictus* (5).

It is a single stranded RNA virus with positive polarity and the genome is 11.8 kb long. The genome is divided into two open reading frames. The 5’ end comprises of all the non-structural proteins (nsPs), while the 3’ end codes for all the structural proteins. The genome is similar to messenger RNA and consists of cap and poly A tail. nsP1 is involved in the formation of viral cap, whereas nsP2 is a crucial protein as it is multifunctional and is an integral component of viral replication complex. On the other hand, nsP3 is the least explored non-structural protein, while nsP4 carries the RNA dependent RNA polymerase activity. In addition to that, there are structural proteins, capsid, 6k and envelope glycoproteins that help in virus particle formation (6).

All the viruses utilize a plethora of strategies to manipulate the host pathways to benefit their own survival and effective replication. One of the strategies is to activate the DDR pathways to utilize the DNA repair proteins. However, the underlying mechanism for the induction of the DNA damage signaling is not completely elucidated till date. It has been reported that several DNA as well as handful of RNA viruses interacts with the cellular proteins involved in DDR which can further lead to up regulation or inhibition of these proteins to facilitate infection (7,8). The DDR pathways consist of a much-orchestrated array of proteins. At the core of the DDR pathways are the members of the phosphoinositide 3 (PI-3) kinase families mainly ataxia-telangiectasia mutated kinase (ATM) which gets activated upon double strand DNA breaks and ATM-Rad3 related kinase (ATR) which is quickly activated in presence of single strand DNA breaks leading to phosphorylation of several downstream protein targets (9).

DNA viruses, such as Papillomaviruses, appeared to activate ATM in differentiating cells and this activation is vital for the formation of virus replication foci (10). Viral DNA replication of Human cytomegalovirus stimulates the ATM pathway and it hampers DNA damage responses by altering the localization of checkpoint proteins to cytoplasm from nucleus (11). Previous investigations have revealed that Herpes simplex virus infection triggers a cellular DNA damage response leading to the activation of ATM and abrogation of ATR signaling pathway that are crucial for viral replication (12,13). In corneal epithelium, Check point kinase-2 (Chk2) plays an important role in HSV-1 replication (14). Whereas, viral interferon regulatory factor 1 interacts with ATM kinase and inhibits the ATM kinase activity in Kaposi’s sarcoma-associated herpesvirus (15). RNA viruses, such as Hepatitis C virus has been found to cause G2/M phase cell cycle arrest (16). Moreover, it is demonstrated that the ATM and Chk2 proteins play pivotal role in RNA replication of this virus. It is also found that HCV-NS5B interacts with both ATM and CHK2, while HCV-NS3-NS4A interacts with ATM (17). Japanese encephalitis virus reduces the proliferation of neural progenitor cells and modulates the cell cycle (18).

As the DDR pathways have never been explored in Alphaviruses, an attempt was made to determine the importance of the DDR pathways in CHIKV infection and to uncover the mechanisms by which CHIKV modulates this. In the current study, it was found that CHIKV infection triggers the activation of ATM/Chk2 and ATR/Chk1 pathways which leads to increased DNA damage and efficient CHIKV infection.

## Materials and Methods

### Cells and Virus

Vero and Human Embryonic Kidney 293T (HEK293T) cells were obtained from NCCS, India. They were cultured in Dulbecco’s modified Eagle’s medium, (DMEM, PAN Biotech, Germany) supplemented with 10% FBS (DMEM, PAN Biotech, Germany), Gentamycin and Penicillin-streptomycin (Sigma, USA). Both the cells were kept at 37 °C and 5% CO2. The Indian CHIKV strain (accession no. EF210157.2), was gifted by Dr. M.M Parida (DRDE, Gwalior).

### Antibodies and inhibitors

The CHIKV-nsP2 antibody, used in this study was developed by our group (19). Chk2, pChk2, Chk1, pChk1 and γH2A.X (Ser-135) antibodies were procured from Cell Signaling Technologies (Cell Signaling Inc, USA). E2 antibody was gifted by Dr. M.M. Parida (DRDE, Gwalior). GAPDH was purchased from abgenex, India. ATM/ATR Kinase Inhibitor (AAKi) was acquired from Sigma Aldrich, USA.

### Alkaline Comet assay

Mock and CHIKV infected cells were harvested at 15hour post infection (hpi), mixed with low melting agar and spread onto microscopic slides precoated with normal agarose. Electrophoresis was performed at 25 volts for 20 min with the agarose gel electrophoresis system (Bio-Rad) and processed for Alkaline Comet assay as previously described (20). The degree of DNA damage was quantified by investigation of randomly selected images in each set (the uninfected and infected groups) with the Casplab software.

### Cytotoxicity Assay

The cytotoxicity of AAKi in Vero cells was determined using EZcount™ MTT cell assay kit (Himedia, India) as per the manufacturer’s protocol. Approximately 20000 number of cells were plated in 96 well plate one day prior to the experiment. The cells with 80% confluency, were treated with different concentrations of inhibitor for 15 hr while DMSO used as a reagent control. The metabolically active cell percentage was compared with the control cells and cellular cytotoxicity was determined as described before (21).

### Viral infection

Vero cells were infected by CHIKV with different multiplicity of infection (MOIs) as per the protocol described before (21). The cytopathic effect (CPE) was seen under microscope (20 X magnification) and the cells as well as supernatants were harvested for various downstream processing.

### Plaque Assay

To calculate viral titre, plaque assay was performed with the mock, infected, and infected with drug treated cell supernatants. For this, after infection cells were overlaid with DMEM containing methylcellulose (Sigma, USA) for 4 days followed by fixed with 8% formaldehyde and stained with crystal violet as described earlier (22).

### Confocal Microscopy

Immunofluorescence analyses were performed as described earlier (23). Briefly, Vero cells were grown in the coverslips, were infected with CHIKV. At 15 hpi cells were fixed with 4% paraformaldehyde and were incubated with primary antibodies of viral proteins nsP2 (1:1000) and host p-Chk2 (1:750) for 1 hr followed by 1X PBS wash. After primary antibodies, incubation was carried out with Alexa Fluor 488 anti-mouse and Alexa Flour 594 anti-rabbit respectively for 45 mins. Then, antifade reagent (Invitrogen) was used to mount coverslips to reduce photobleaching. The Leica TCS SP5 confocal microscope (Leica Microsystems, Germany) was used for acquiring images.

### Animal Studies

Animal experiments were executed in strict compliance with the Committee for the Purpose of Control and Supervision of Experiments on Animals (CPCSEA) of India. All procedures and tests were reviewed and permitted by the Institutional Animal Ethics Committee (ILS/IAEC-246-AH/SEPT-21). 10-12 days old mice were infected subcutaneously with 10^7 PFU of CHIKV at the flank region of the hind limb and mock mice with serum free media as previously described (23). AAKi (2mg/kg) was administered orally to the treated group of mice (n=3) at every 24 hours interval up to 4 days post infection (dpi). Solvent was given to mock and Infection-control group (n=3) of mice. Disease symptoms were observed daily. At 5 dpi, mice were sacrificed. From the blood, serum was isolated to assess viral load. Tissues were collected for Western blot and immunohistochemistry analyses.

### qRT-PCR

Cells were harvested and RNA extraction was performed using TRIzol (Invitrogen, US). An equal volume of RNA was used for the cDNA synthesis by using the First Strand cDNA synthesis kit (Invitrogen, US) and CHIKV specific primers for the E1 gene was used for qRT-PCR. Same amount of supernatants or serum samples was used for viral RNA extraction using the QIAamp viral RNA minikit (Qiagen, Germany) as per the manufacturer’s instructions. The RNA copy number was obtained with the Ct values plotted against the standard curve (23).

### siRNA transfection

Gene silencing of Chk2 and Chk1 was performed by using siRNA as previously described (22). In brief, monolayers of HEK293T cells with 70% confluency were seeded in 6 well plate and was transfected with Lipofectamine 2000 (Thermo Fisher Scientific, USA) using various concentrations of siRNA (30, 60 and 90 pm) corresponding to Chk2 mRNA sequence (sense strand GAAGAGGACUGUCUUAUAAdTdT, CUCAGGAACUCUAUUCUAUdTdT, GAUCUUCUGUCUGAGGAAAdTdT) and Chk1 mRNA sequence (sense stand GAACCAGUUGAUGUUUGGU dTdT, CUCACAGGGAUAUUAAACC dTdT, UGUGUGGUACUUUACCAUA dTdT) or with siRNA negative control. Twenty-four hr after transfection, they were infected with CHIKV with MOI 0.1. The supernatants were collected after 15 hpi and subjected to Plaque assay while cells were harvested for total RNA isolation and Western blot analysis.

### Coimmunoprecipitation

Mock and CHIKV infected Vero cells were harvested at 6hpi followed by lysis with RIPA buffer and subjected to coimmunoprecipitation by Dynabeads® Protein A Immunoprecipitation Kit (Thermo Fisher Scientific, USA) as described earlier (24).

### Protein-protein interaction

The ClusPro2.0 webserver (25) was used to study the protein-protein interaction following reported methods (24). Briefly, the structures of CHIKV nsP2 (PDB ID: 3TRK), Chk1 (PDB ID: 4FSN) and Chk2 (PDB ID: 3I6U) were extracted from the protein data bank (PDB) database. The structures were processed and energy minimised using the Discovery Studio program. The structures of CHIKV nsP2 was used as ligand for the protein-protein interaction study in the ClusPro2.0 webserver. From the four different outputs, the “balanced” outputs were used for further analysis (24). The first docking solution clusters usually have the largest members. Thus, this was taken for further visualization in the PyMol software. The most stable complex was further evaluated using the HyPPI (ProteinPlus) web server (26). Based on the energy (hydrophobic effect) emerging due to binding of protein subunits and the quotient of interface area ratios this server classifies protein-protein complex as permanent, transient and crystal artefact (26,27). The interaction complex was subjected to this analysis to understand the probable nature of the interaction.

### Western blot

Western blot analysis was executed as earlier described (23). In brief, mock, virus infected and drug treated cells were harvested at different hpi according to the experiments and lysed subsequently with equal volume of RIPA buffer. Snap frozen mock, infected and infected-drug treated mice tissues were homogenised by hand homogeniser and lysed in RIPA buffer using the syringe lysis method. Proteins were separated into a 10% SDS-polyacrylamide gel and was transferred onto PVDF membrane. Membrane was probed with antibodies as recommended by the manufacturer. Blots were developed by Immobilon Western Chemiluminescent HRP substrate (Millipore, US). The Image J software was used for quantification of protein bands from three independent experiments.

### Immunohistochemistry

For Immunohistochemistry studies, formalin-fixed tissue samples were dehydrated and embedded in paraffin wax, and serial paraffin sections of 5 mM were obtained as described previously (23). Briefly, tissue sections were incubated with primary E2 antibody followed by secondary Alexa Fluor 594 antibody (anti-mouse; Invitrogen, US). The slides were mounted with DAPI (Invitrogen, US), and coverslips were applied to the slides.

### Cell cycle analysis

Mock and infected cells were washed, harvested and resuspended in chilled 70% ethanol for fixation. Further it was stained with propidium iodide staining solution containing RNaseA and cell cycle analysis was performed using LSR Fortessa as previously described (28).

### Isolation and CHIKV infection in human peripheral blood mononuclear cells (hPBMCs) and flow cytometric staining

Human PBMCs were isolated from the blood samples of three healthy donors as previously described (23). All the adherent cells were detached after 5 days. MTT assay was performed with various concentrations of AAKi for the adherent hPBMCs. After 1 d, adherent cells were pretreated with AAKi at 25 μM for 2 h and subjected to CHIKV infection at a MOI of 5 for 2 h. Infected cells were harvested at 12 hpi followed by fixation with 4% paraformaldehyde (PFA). Next, the cells were subjected to intracellular staining to detect viral protein E2 and surface staining for immunophenotyping of adherent population using flow cytometry. For immunophenotyping, fluorochrome conjugated anti-human CD3, CD11b, CD14, and CD19 antibodies (Abgenex, India) were used as previously described (23).

### Statistical analysis

Data are shown as SEM for three independent experiments. The GraphPad Prism version 8.0.1 software was used for statistical analyses. For two group’s comparison, unpaired two-tailed Student’s t-test was performed. For three or more groups, the one-way or two -way ANOVA with Bonferroni post-test was used. P values are specified at each figure or at their respective legend, and figure symbols represent *p < 0.05, **p < 0.01, ***p < 0.001, ****p < 0.0001.

## Results

### Induction of DNA damage signalling pathways following CHIKV infection

Chk2 and Chk1 are protein kinases, which are downstream mediator proteins of ATM and ATR arm of DDR pathways (9). To investigate whether DNA damage response pathway is activated during CHIKV infection, the phosphorylation status of Chk2 and Chk1 in CHIKV infected cells were examined. First, Vero cells were infected with CHIKV with MOI 2 and harvested at different time points and protein levels were evaluated by Western blot. It was found that Chk2 and Chk1 proteins were phosphorylated in CHIKV infected cells (Figure 1A and B). Interestingly, the levels of phosphorylation of both Chk2 and Chk1 protein expression were enhanced gradually with increasing time points in CHIKV infected cells as compared to the corresponding mock cells. The total Chk2 and Chk1 protein levels remain unaltered in every time point as shown in Figure 1A and B. To further validate that a DNA damage response was induced by CHIKV, phosphorylation of the γH2A.X was examined. Vero cells were infected with CHIKV and processed for Western blot analysis. It was found that γH2A.X was phosphorylated following CHIKV infection as compared to mock cells (Figure 1C). To further confirm phosphorylation of p-Chk2 following CHIKV infection, Immunofluorescence analyses was performed and p-Chk2 was found to be colocalizing with nsP2 protein of CHIKV with a Pearson’s Coefficient: r = 0.691 as shown in figure 1D. Together, these findings indicate that DNA damage signaling pathways are activated following CHIKV infection.

**Figure 1.**
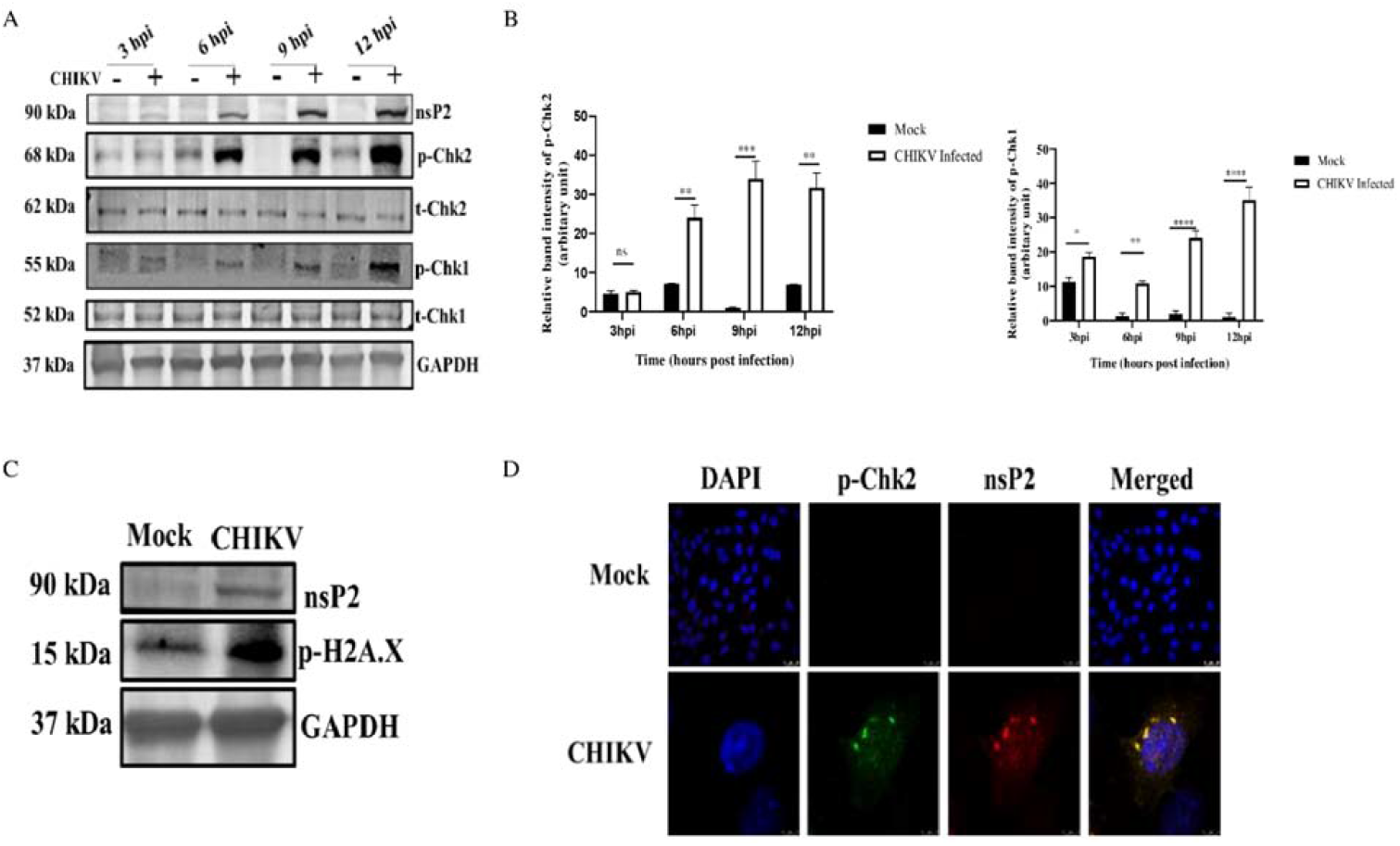
CHIKV infection induces DDR pathways. Vero cells were mock or infected with CHIKV and harvested at various time points. (A) Western blot was performed using nsP2, p-Chk.2 (Thr-68), total Chk2, p-Chk1(Ser354), total Chk1, and GAPDH antibodies. **(B)** Bar diagram showing relative band intensities of p-Chk2 and p-Chk1 at different time post infection. Data of three independent experiments are shown as mean ± SEM. (C) Western blot depicting the level of p-H2A.X (Ser-139) in mock and infected samples. **(D)** Mock or infected Vero cells were stained with nsP2 and p-Chk2 antibodies. Nuclei were counterstained with DAPI.

### DNA Damage is enhanced in CHIKV infected cells

To examine whether DNA damage was induced during CHIKV infection, alkaline comet assay was performed. It was found that, no DNA migration occur among most of the control cells while increase in the length of DNA migration was observed in CHIKV infected cells (Figure 2A and B). The extent of DNA damage in CHIKV infected cells was assessed by the tail moment using the Casplab software. CHIKV infection enhanced the degree of DNA damage by 95% compared to that in mock cells, indicating good amount of DNA damage (Figure 2A and B). Taken together, the findings demonstrate that DNA damage is induced upon CHIKV infection.

**Figure 2.**
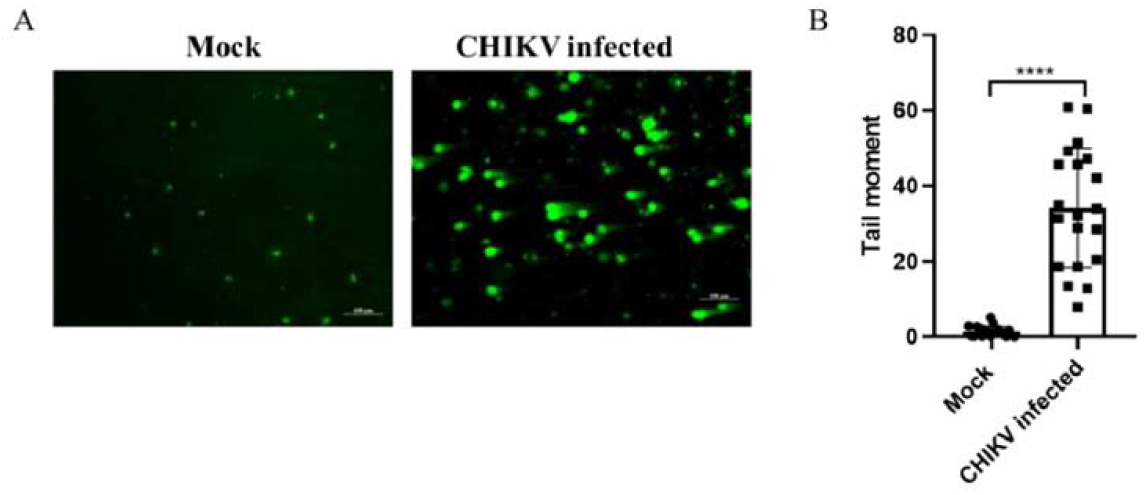
DNA damage is induced by CHIKV infection. Mock and CHIKV infected Vero cells were harvested at 18 hpi. Alkaline Comet assays were performed. (A) Image depicting the DNA damage in CHIKV infected and mock cells (B) The cluster plot analysis portraying tail moment in mock and CHIKV infected samples. Data of three independent experiments are shown as mean ± SEM.

### Both ATM-ATR pathways facilitate efficient CHIKV infection

In order to understand the importance of DDR pathways on CHIKV infection, AAKi was used. To determine the cytotoxicity of the inhibitor in Vero cells, the MTT assay was performed. It was observed that 98% cells were viable at all the concentrations of the drug (Figure 3A). After CHIKV infection and drug treatment (10, 20 and 30μM) cytopathic effect was observed under microscope. Clear reduction in cytopathic effect (CPE) was found as compared to the DMSO control (Figure 3B). This result was further confirmed by Western blot analysis where 2, 8.5 and 150 -fold reduction was noted in nsP2 level, in presence of 10, 20 and 30μM concentrations of the drug respectively (Figure 3 C and D). Similarly, it was observed that there was 50-fold reduction in the level of E1 gene by RT-PCR as shown in figure 3E. Next, plaque assay was performed from the cell culture supernatants and interestingly, it was observed that the viral titers were also reduced by 3, 15.8 and 475-fold in presence of 10, 20 and 30μM concentrations of the drug respectively (Figure 3F). Collectively, these results indicate that both ATM as well as ATR pathways are critical for efficient CHIKV infection.

**Figure 3.**
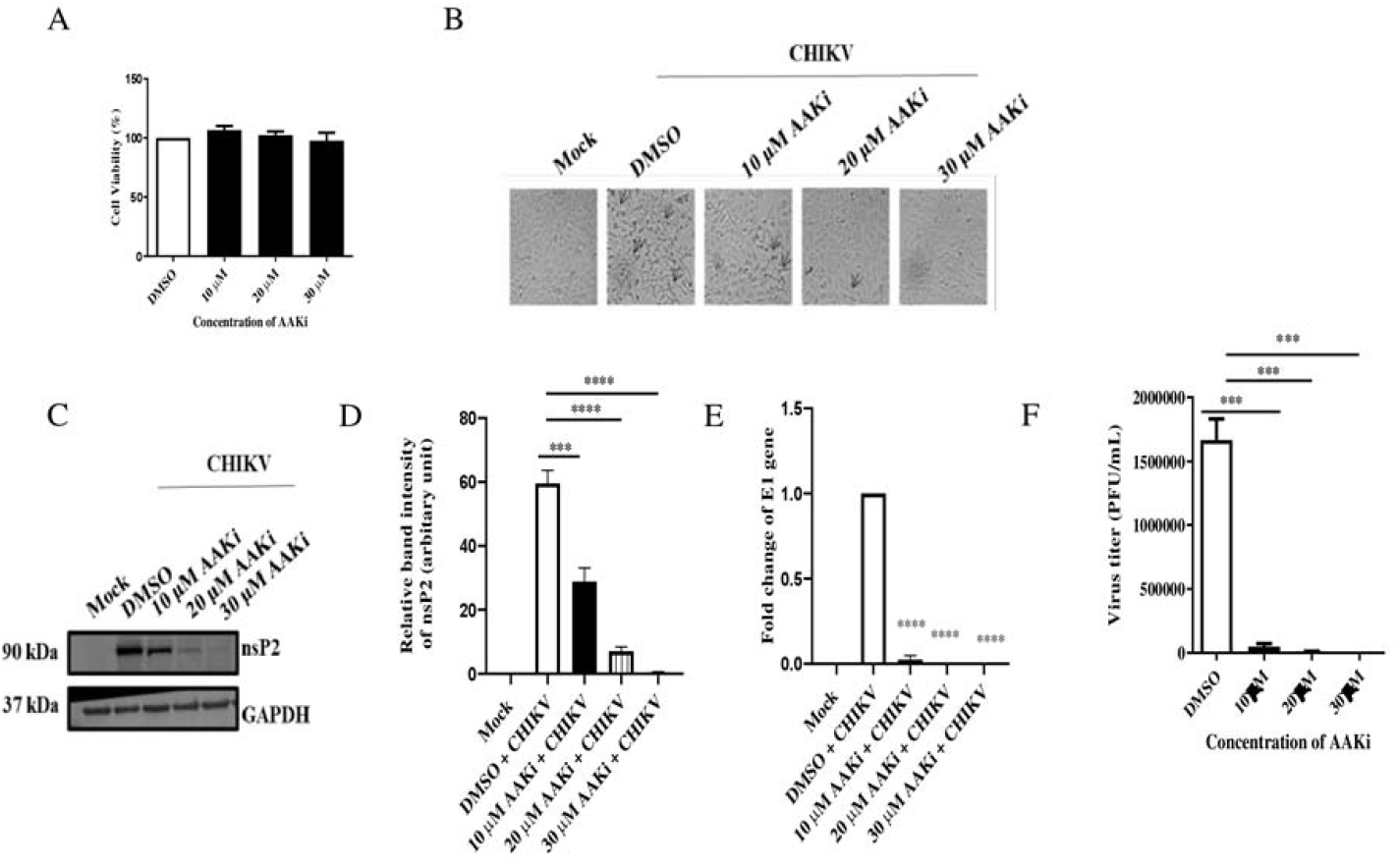
AAKi inhibits CHIKV infection. (A) Vero cells were treated with different concentrations (10, 20, and 30μM) of AAKi for 15 h and the cytotoxicity of the cells were estimated by MTT assay. (A) Bar diagram showing the viability of cells. (B) image depicting the CPE of cells which was observed under microscope (Magnification -10X) for cytotoxicity with different concentration (10, 20, and 30μM) of the drug. (C) Western blot image of mock, infected and infected plus treated cells. GAPDH served as the loading control. (D) The relative band intensity of nsP2 is shown as bar diagram. (E) Whole RNA was isolated from the mock, CHIKV-infected and drug-treated cells, and the CHIKV E1 gene was amplified by qRT-PCR. Bar diagram displaying the fold change of viral RNA. (F) Bar diagram representing the viral titer in the cell supernatant. Data of three independent experiments are shown as mean ± SEM.

### Efficient inhibition of CHIKV infection by AAKi in mice

To measure the antiviral effect of AAKi *in vivo*, 10-12 days old C57/BL6 mice were infected with CHIKV and treated with 2mg/kg AAKi at every 24 hours interval up to 5 dpi. The infected mice showed loss of body weight, paralysis in limbs and weakened mobility, whereas, AAKi -treated mice showed no such anomalies (Figure 4A and B). Viral RNA was isolated from the pooled serum samples (from respective group) and RT-PCR was carried out to estimate the CHIKV copy number and it was found that the viral load was reduced to 71 % in AAKi -treated mice in comparison to control (Figure 4C). Moreover, Western blot analysis was also executed to determine the viral protein level in infected and treated group of mice tissues. 95% and 58 % reduction of nsP2 protein level in muscle and brain was observed in the treated mice (Figure 4D and E). Further, immunohistochemistry analysis revealed the decrease of E2 level in CHIKV infected muscle upon AAKi treatment (Figure 4F) These results indicate that AAKi can protect mice against CHIKV infection.

**Figure 4.**
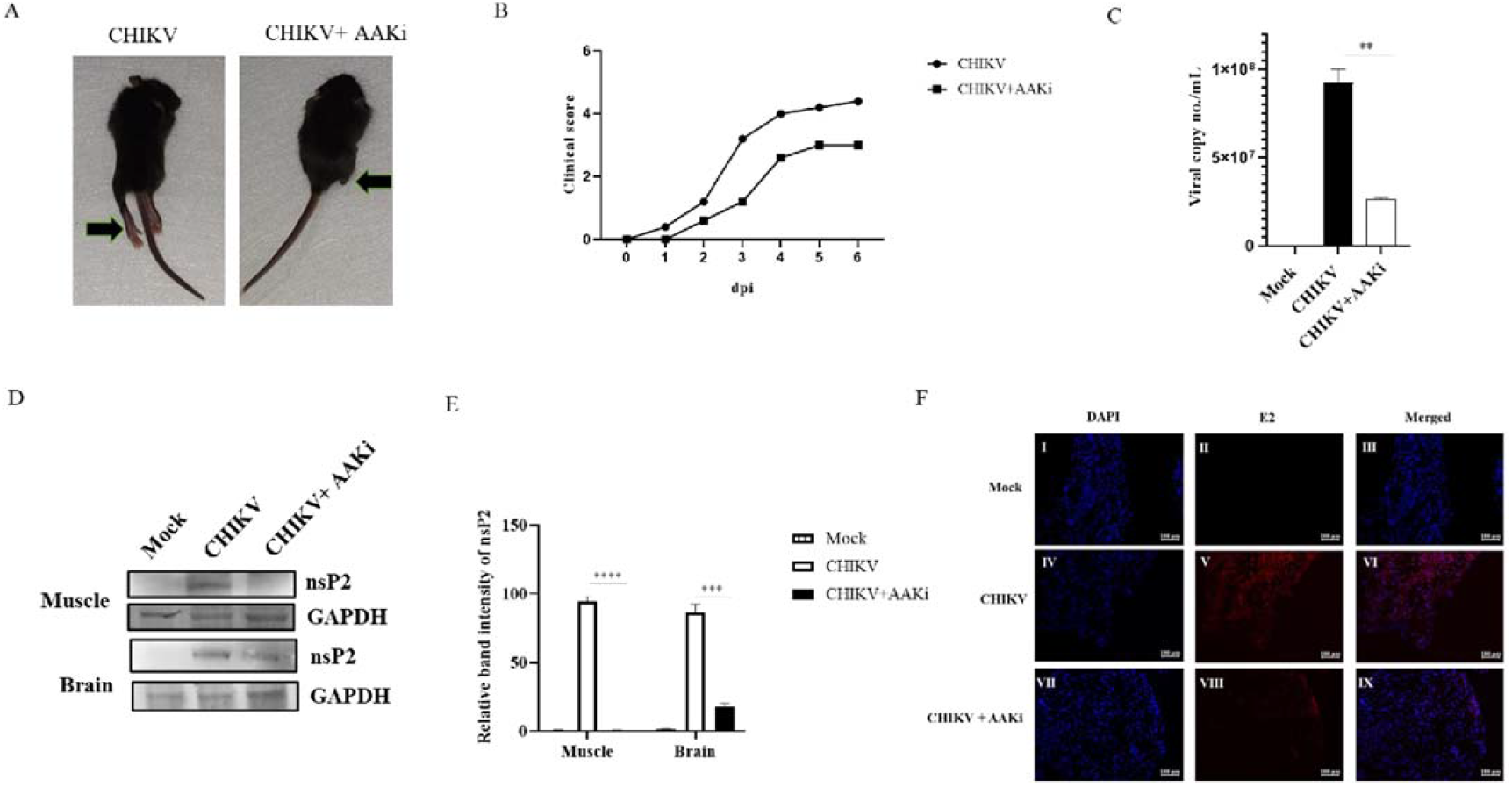
Efficient inhibition of CHIKV infection by AAKi in mice. C57BL/6 mice were infected subcutaneously with 10^7^ PFU of CHIKV and treated with 2 mg/kg of AAKi at 24 h intervals up to 4 dpi. Mice were sacrificed at 5 dpi, and serum and different tissues were collected for experiments. An equal volume of sera was taken to isolate viral RNA. cDNA was synthesized, and the E1 gene was amplified by qRT-PCR to determine the viral RNA copy number. (A) Image of CHIKV-infected and drug-treated mice. (B) Graph showing clinical score of the disease symptoms during CHIKV infection which were monitored from 1 dpi to 5dpi. (C) Bar diagram showing CHIKV RNA copy number/ml in virus-infected and drug-treated mice serum. (D) Western blot showing the viral nsP2 protein in muscle and brain tissue samples. GAPDH was used as a loading control. (E) Bar diagram showing the relative band intensities of nsP2 in muscle and brain tissue samples from infected or drug-treated animals. (F) Confocal microscopic image panels showing the CHIKV E2-stained muscles from infected or drug-treated mice. Data of three independent experiments are shown as mean ± SEM.

### Chk2 is crucial for CHIKV replication

To investigate the importance of Chk2 protein for CHIKV infection, 30 and 60 pm concentrations of siRNA was used to silence the expression of Chk2 gene in HEK293T cells. Cells were harvested at 24 hpt and processed for Western blot to measure Chk2 level. It was observed that Chk2 protein level was reduced by 62.5% in 60 pm concentration of siRNA as compared to control (Figure5A and B). Thus, 60 pm concentration of Chk2 siRNA was selected for further experiments. Next, the siRNA transfected cells were infected with CHIKV and supernatants and the cell lysates were collected at15hpi were processed for plaque assay and Western blot respectively. Interestingly, it was found that there was 73.7% reduction in the viral progeny formation in comparison to the control (Figure 5C). Similarly, there was 50% reduction in the level of E1 gene by RT-PCR as shown in figure 5D. While, in Western blot analysis, it was observed that the nsP2 protein level was reduced by 88.3% after siRNA transfection (Figure 5E and F). Chk2 level was also less as expected (Figure 5E and G) and together the results suggest that Chk2 is a crucial host factor for CHIKV infection.

**Figure 5.**
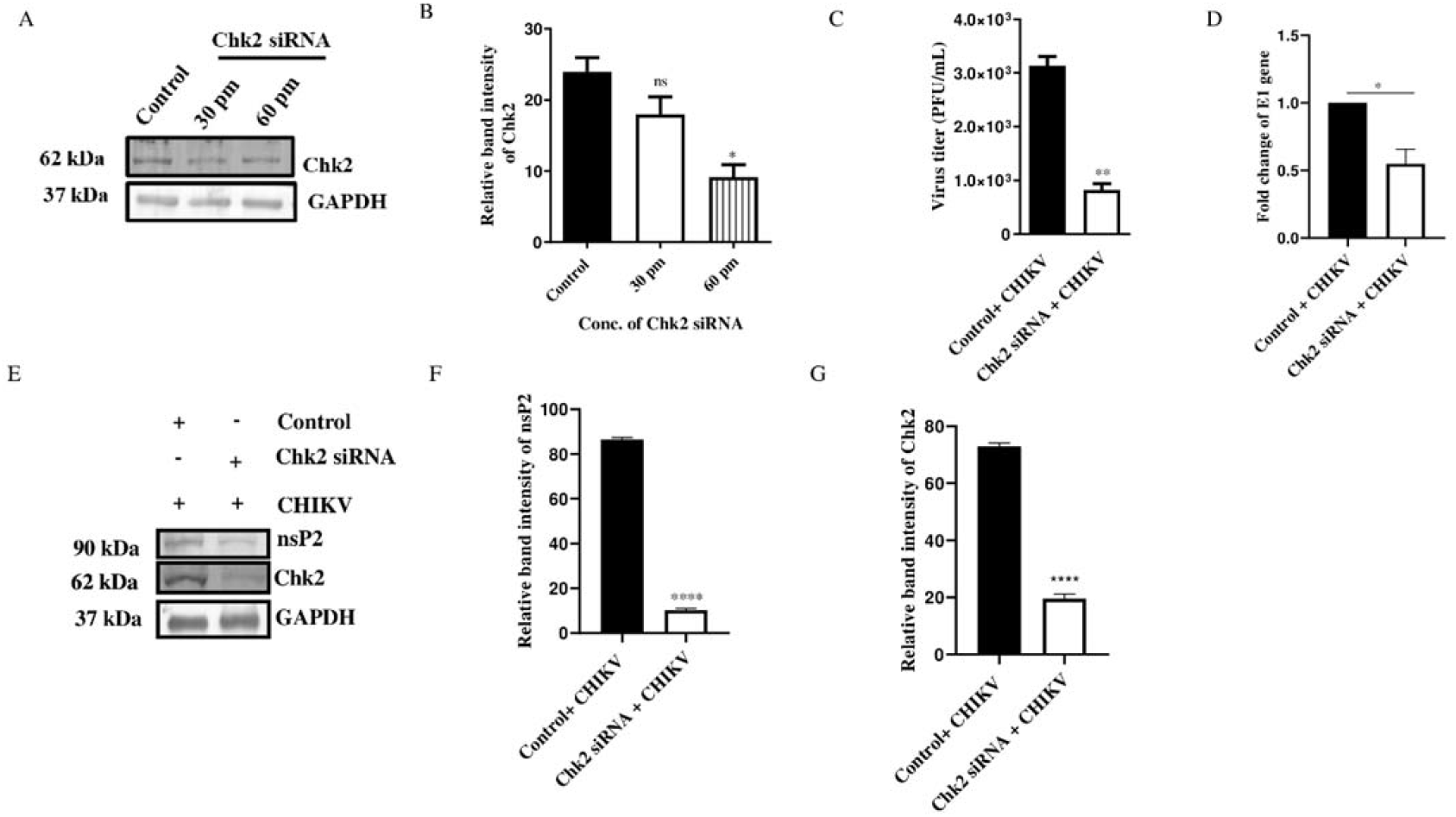
Chk2 is essential for CHIKV infection. HEK293T cells were mock transfected or transfected with 30, 60 and 90 pm of Chk2 siRNA. (A) Chk2 level was estimated in Western blot and GAPDH was used as loading control. (B) Bar diagram showing relative band intensity of Chk2 protein. After 24 hpt (60 pm), cells were super infected with CHIKV (MOI of 0.1) and harvested at 15 hpi for further downstream experiments. (C) Bar diagram representing the viral titer in the cell supernatant of control+CHIKV and Chk2 siRNA+CHIKV samples. (D) Bar diagram showing the fold change of E1 gene of control+CHIKV and Chk2 siRNA+CHIKV samples. (E) Western blot showing the nsP2 and Chk2 protein levels after transfection and super infection with CHIKV. (F) and (G) Bar diagrams depicting relative band intensities of the nsP2 and Chk2 proteins. Data of three independent experiments are shown as mean ± SEM.

### Chk1 is essential for CHIKV replication

In order to study the role of Chk1 in CHIKV infection, 30 pm concentration of siRNA was used to silence the expression of Chk1 gene in HEK293T cells. After 24 hpt. Cells were harvested at 24 hpt and processed for Western blot. It was observed that Chk1 protein level was reduced by 59.3% using 30 pm concentration of siRNA as compared to control (Figure 6A and B). Next, the siRNA transfected cells were infected with CHIKV and supernatants and the cell lysates collected at 15hpi were processed for plaque assay and Western blot. Interestingly, it was found that there was 78.8% reduction in the viral progeny (Figure 6C). While more than 50% reduction was observed in the level of E1 gene by RT-PCR as shown in figure 6D. Moreover, in Western blot analysis, it was observed that the nsP2 level was reduced by 79% after siRNA transfection (Figure 6E and F). Chk1 level was also reduced as expected (Figure 6E and G) and the results suggest that Chk1 protein is essential for CHIKV infection.

**Figure 6.**
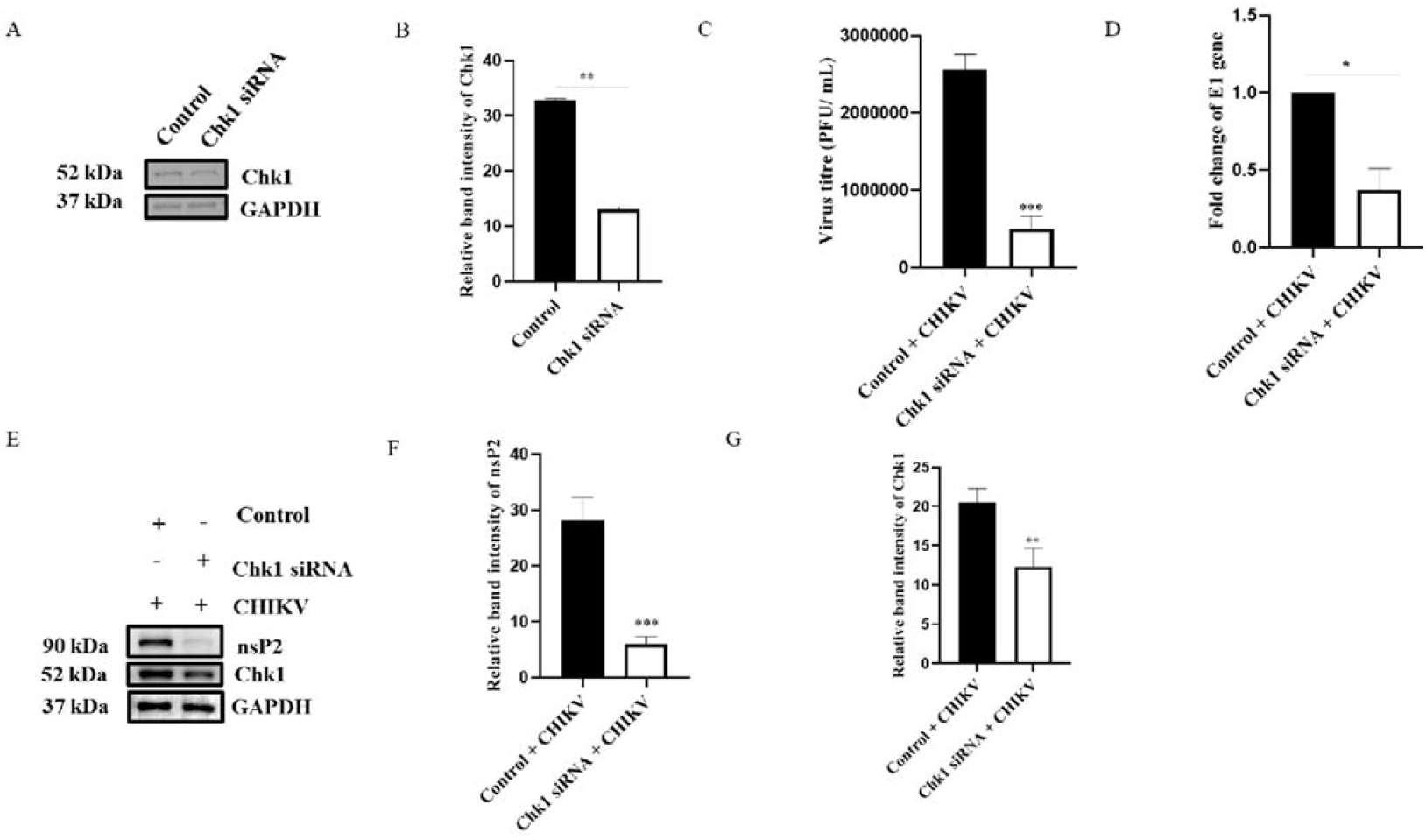
Chk1 is crucial for CHIKV infection. HEK293T cells were mock transfected or transfected with 30 pm of Chk1 siRNA. (A) Chk1 level was estimated in Western blot and GAPDH was used as loading control. (B) Bar diagram showing relative band intensity of Chk1 protein. After 24 hpt (30 pm), cells were super infected with CHIKV (MOI of 0.1) and harvested at 15 hpi for further downstream experiments. (C) Bar diagram representing the viral titer in the cell supernatant of control+CHIKV and Chk1 siRNA+CHIKV samples. (D) Bar diagram showing the fold change of E1 gene of control+CHIKV and Chk1 siRNA+CHIKV samples. (E) Western blot showing the nsP2 and Chk1 protein levels after transfection and super infection with CHIKV. (F) and (G) Bar diagrams depicting relative band intensities of the nsP2 and Chk1 proteins. Data of three independent experiments are shown as mean ± SEM.

### CHIKV-nsP2 interacts with both Chk1 and Chk2 during CHIKV infection

CHIKV infection altered the phosphorylation of host Chk2 and Chk1 proteins, hence their association with the nsP2 protein was investigated. CHIKV infected cells were harvested at 6 hpi and processed for co-immunoprecipitation followed by Western blot analysis. It was found that the CHIKV-nsP2 protein was pulled by both the Chk2 and Chk1 proteins (Figure 7A and B). This result suggests that CHIKV-nsP2 interacts with Chk2 and Chk1 during CHIKV infection.

**Figure7.**
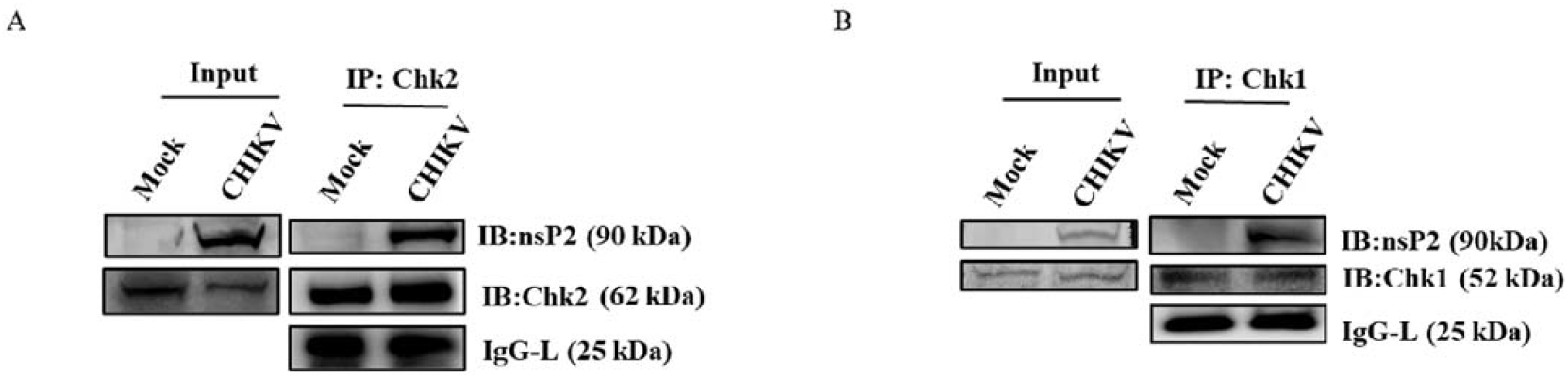
CHIKV-nsP2 interacts with the host Chk2 and Chk1 proteins. Vero cells were infected with CHIKV for 6 hpi. The cell lysates were co-immunoprecipitated with the Chk2 and Chk1 antibodies respectively. (A) Western blot analysis depicting the expressions of nsP2 and Chk2 in the whole cell lysate (left), co-immunoprecipitation analysis showing the interaction of the CHIKV nsP2 and Chk2 protein (right). (B) Western blot analysis depicting the expressions of nsP2 and Chk1 in the whole cell lysate (left), co-immunoprecipitation analysis showing the interaction of the CHIKV nsP2 and Chk1protein (right).

### Identification of the crucial amino acids responsible for the nsP2-Chk2 and nsP2-Chk1 interactions

In order to unravel the amino acid residues responsible for the interaction of nsP2-Chk2 and Chk1, protein-protein docking was carried out as mentioned above. As the balanced outputs of the protein-protein docking studies takes into account all possible modes of interaction, the most stable conformation of these outputs was analysed using the PyMol software. Close interaction was predicted between Chk1 (PDB ID: 4FSN) and nsP2 (PDB ID: 3TRK) as shown in Figure 8A and B. Arg, 1105, Leu1103, Arg 1278, Arg 1284, Arg 1308 and Asn1317 of the nsP2 was involved in multiple polar interactions with Glu7, Asp8, Glu134, Asp94, Gly18 and Gln13 of the Chk1 respectively (Table S1). Further evaluation of the interaction complex by the ProteinPlus web server revealed that the complex is not a mere artifact and has relatively higher score (65%) as a permanent complex than a transient complex (24%). Relatively, less number of interactions were predicted for interaction between nsP2 (PDB ID: 3TRK) and Chk2 (PDB ID: 3I6U) (Table S2). Nonetheless, strong polar interactions (H bond length<2Å) were observed between Asn-1082, Thr-1200, Arg-1195, Asp-1198, Pro-1182 of nsP2 and Lys-241, Arg-240, Ser-210, Lys-224, Asn-186 of Chk2 respectively (Figure 8C and D). It showed higher score (85%) as a transient complex than a permanent complex (13%). Thus, the protein-protein docking analysis identifies the crucial amino acids responsible for the interaction between CHIKV-nsP2 and Chk1 as well as Chk2.

**Figure 8.**
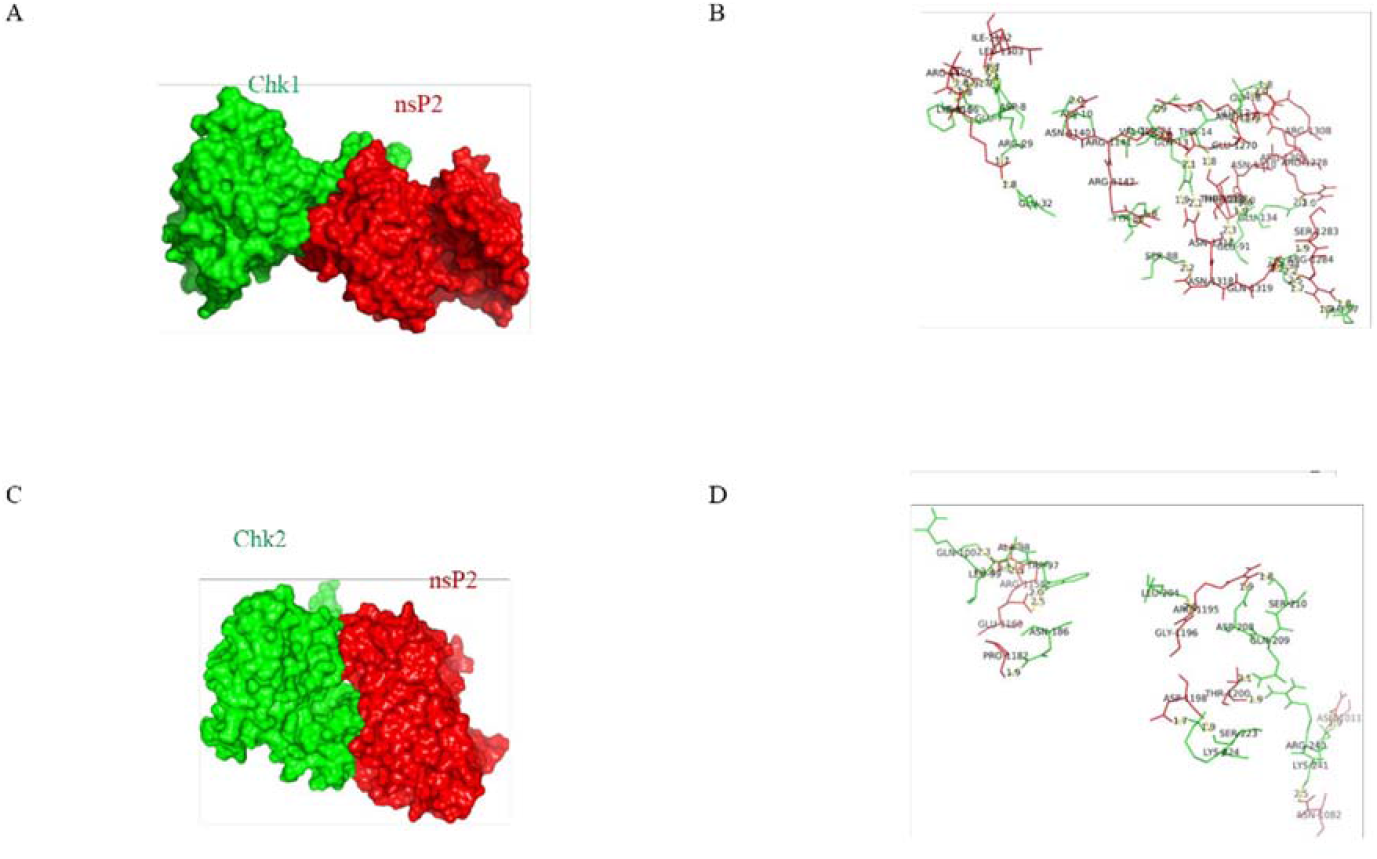
Protein-protein docking analysis shows the probable interaction of CHIKV-nsP2 with host Chk1 and Chk2. The protein-protein docking was performed using the ClusPro 2.0 web server. (A) Model showing the probable interaction of nsP2 (red surface) with host Chk1 (green ribbon). (B) Figure highlights polar interactions (yellow bridge) between residues of nsP2 (red) and host Chk1 (green). (C) Model showing probable interaction of nsP2 (red surface) with host Chk2 (green ribbon). (D) Figure depicts polar interactions (yellow bridge) between residues of nsP2 (red) and Chk2 (green).

### CHIKV infection leads to both G1 and G2 arrest

Viruses are known to cause cell cycle turmoil (28). Here, it was found that both the Chk2 and Chk1 proteins were modulated upon CHIKV infection, hence it was interesting to investigate the cell cycle arrest. This was analysed by measuring DNA content with PI staining for mock- and CHIKV-infected cells at 18hpi. It was observed that CHIKV infection augmented the percentage of cells in the G1 phase from 58.46% to 74.87% whereas, in the G2 phase it was increased from 5% to 18% in Vero cells (Figure 9), indicating that CHIKV infection induces cell cycle arrest in both the G1 as well as G2 phases.

**Figure 9.**
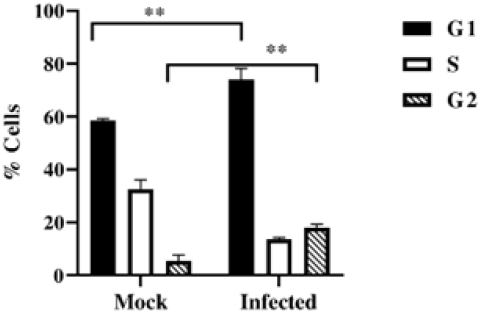
Cells are arrested in G1 and G2 phases following CHIKV infection. Vero cells were mock infected and infected with CHIKV and cells were collected at 15 hpi. Cell cycle analysis was performed with PI staining. Bar diagram showing quantitation of the data. Data of three independent experiments are shown as mean ± SEM.

### AAKi abrogates CHIKV infection in hPBMC

Immunological characterization of hPBMC derived adherent population was carried out using antibodies against specific markers of B cell (CD19), T cell (CD3), monocyte-macrophage cells (CD11b and CD14) and analyzed by flow cytometry (Figure 10A). It was found that the adherent population is highly enriched with CD14+CD11b+ monocyte-macrophage cells. Then adherent populations were scraped out and plated in 96 well plates at a confluency of 5000 cells per well for MTT assay. Various concentrations of AAKi (1, 2.5, 5, 7.5, 10, 25 and 50 μM) were added after 24 h of seeding and cells were incubated for 24 h at 37° C in dark. 25 μM AAKi concentration was found to show cell viability of more than 95% (Figure 10B), hence all the experiments were further performed at 25 μM concentration of AAKi. Surprisingly, it was found that there was 97% reduction in the viral progeny formation in comparison to the control as shown in figure 10C. Next, the effect of AAKi in hPBMC derived adherent populations of 3 healthy donors were investigated upon CHIKV infection *ex vivo*. CHIKV infected populations showed 27.93 ± 0.9 % E2 positive cells, whereas pre-incubation with 25 μM of AAKi for 3h lead to a decrease in E2 positive cells upto 0.86± 0.4 % (Figure 10D and E). Taken together, the data suggest that AAKi can abrogate CHIKV infection significantly in human PBMC derived monocyte-macrophage populations *ex vivo*.

**Figure 10.**
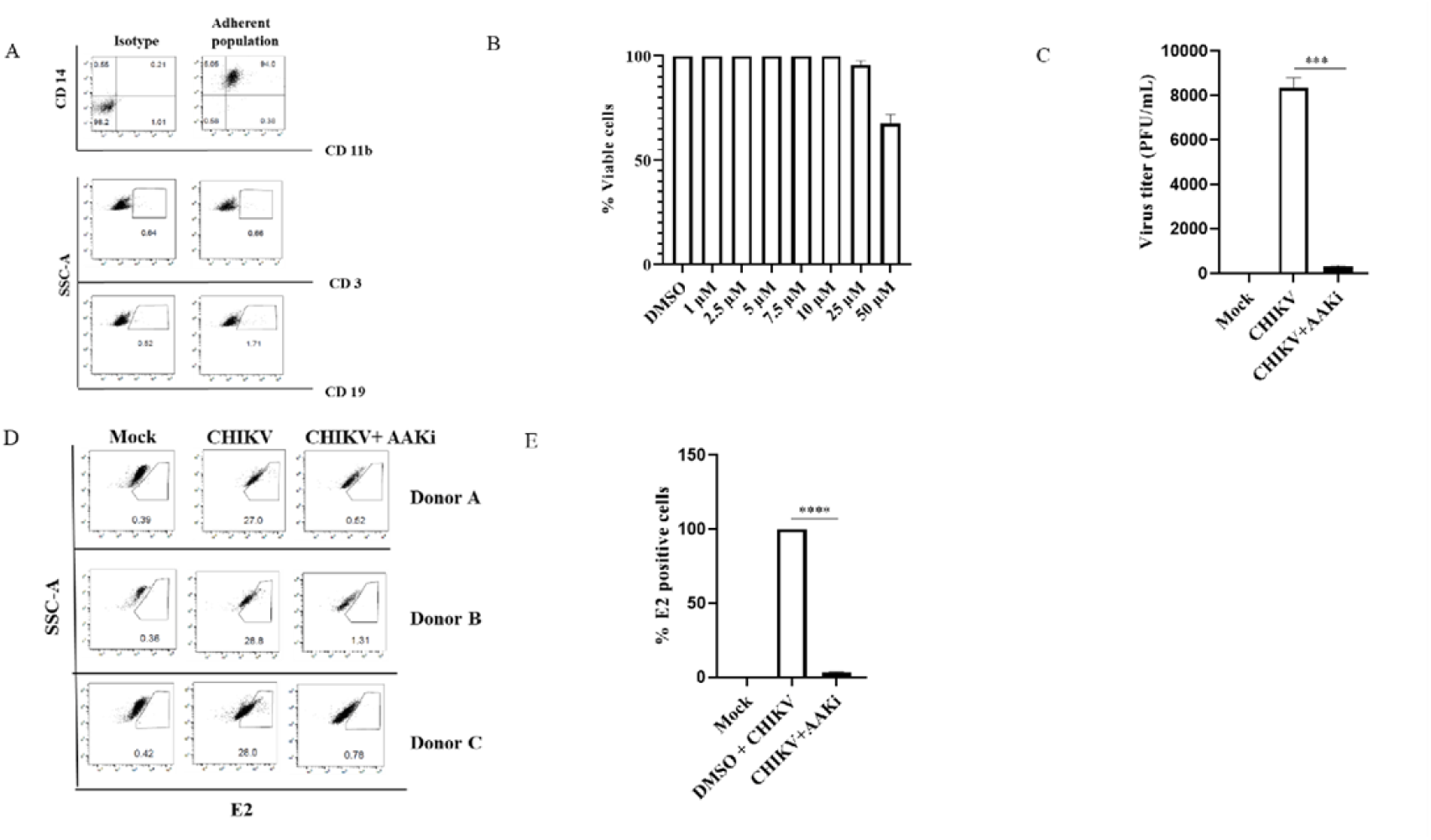
AAKi reduces CHIKV infection in hPBMC-derived monocyte-macrophage populations *ex vivo*. Human PBMCs were isolated from the blood samples of three healthy donors and infected with CHIKV. (A) Dot plot showing the percentages of B cells (CD19), T cells (CD3), and CD141CD11b1 monocyte-macrophage cells from adherent hPBMCs by flow cytometry. (B) Bar diagram representing the cytotoxicity of AAKi in hPBMC-derived adherent cell populations by MTT assay. (C) Bar diagram depicting percentage of the viral particle formation obtained by plaque assay. (D) Dot plot showing the percentage of viral E2-positive hPBMC-derived monocyte-macrophage population in mock, CHIKV-infected, and AAKi treated CHIKV infection from three healthy donors’ hPBMCs obtained using flow cytometry. (E) Bar diagram showing the percentage of positive cells for CHIKV E2 protein, as derived by flow cytometry assay.

## Discussion

Viruses being intra-cellular parasite, need several host cell machineries so as to achieve effective replication of their own genome, along with virus-encoded enzymes. One of the strategies is to hijack the DDR pathways. Several DNA as well as handful of RNA viruses interact with the cellular proteins involved in DDR pathways, however, reports with respect to the association of Chk2 and Chk1 in alphavirus infection are scanty. Hence, this study is amongst the first to report that modulation of DDR pathways is crucial for effective CHIKV infection. This work showed for the first time the mechanistic insight of the induction of DDR pathways by CHIKV and interaction of CHIKV-nsP2 with two crucial host factors, Chk2 and Chk1 for efficient viral infection. This information might contribute to the development of effective therapeutics for the control of the CHIKV infection in future.

The study reveals that CHIKV infection activates the Chk2 and Chk1 proteins associated with DDR signaling pathways, as well as increases DNA damage by 95%. Inhibition of both ATM-ATR kinases by AAKi showed that the viral particle formation was reduced by 15.8-fold in presence of 20 μM concentration of the molecule as compared to the control. Additionally, the treatment of mice with this drug has reduced the disease score substantially in CHIKV-infected C57BL/6 mice with 71% decrement in the viral copy number and diminished viral protein level. Gene silencing of Chk2 and Chk1 reduced viral progeny formation around 73.7% and 78% respectively, validating that both Chk2 and Chk1 are crucial host factors for CHIKV infection. Moreover, it was demonstrated that CHIKV-nsP2 interacts with Chk2 and Chk1 during CHIKV infection and protein-protein docking analysis depicted the specific amino acids responsible for these interactions. Further, the data indicated that CHIKV infection induces cell cycle arrest in both G1 as well as G2 phases. Besides, the abrogation of CHIKV infection was also established *ex vivo* in hPBMC-derived monocyte-macrophage populations by AAKi, confirming the importance of DDR pathway for CHIKV life cycle.

In this study, CHIKV has been found to activate the proteins associated with DDR pathways and similar observations have been noted in few other viruses like Bocavirus, Human Parvovirus and BK Polyomavirus (29–32). Here it was also observed that CHIKV phosphorylates γH2A.X which is a hallmark of the DDR induction and that was also reported previously in case of Rift Valley Fever virus (33). The studies have also enlightened about the damage of DNA along with the induction of DDR in few cases like ZIKV infection and MDV infection (20,34) and surprisingly CHIKV also belong to the same category showing DNA damage after viral infection. However, several (RVFV and BK Polyomavirus) viral infection (33,35) do not lead to DNA damage. The actual mechanistic details to understand the role of these host factors for virus infection is not very clear and that requires further investigation.

The findings suggest that the components of DDR pathways are recruited to the nucleus, while CHIKV replication takes place in the cytoplasm only. This raised a question about the modulation of the DDR pathways by the virus to facilitate its replication. Interestingly, the current study exhibited that CHIKV-nsP2 colocalizes with pChk2 in the nucleus and the interaction between Chk1, Chk2 and nsP2 was confirmed by coimmunoprecipitation also. It was also found that HCV encoded NS5B interacts with both ATM and CHK2, while HCV NS3-NS4A interacts with ATM (17). Kaposi’s sarcoma-associated herpesvirus encoded vIRF1 interacts with endogenous ATM (15). ATR and pChk1 were observed to colocalize with LT nuclear foci in Merkel cell polyomavirus (36) indicating that there are viral proteins which can localize in the nucleus to hijack the host machineries for their benefit. The nsP2, Chk2 and Chk1 interactions were further supported by the protein-protein docking analysis that showed involvement of residues in the polar interaction at the protein surface. The complex formed between Chk1 (PDB ID: 4FSN) and nsP2 (PDB ID: 3TRK) showed strong polar interaction (≈37 H bonds) between several residues of CHIKV nsP2 and Chk1. Although strong polar interactions were observed in the CHIKV nsP2-Chk2 complex, the number of polar interactions (≈16 H bonds) were relatively less. On account of higher degree of interaction at the surface, the Protein Plus server classifies the CHIKV nsP2 and Chk1 interaction as permanent complex suggesting higher lifetime of this interaction. In contrast, the interaction complex of CHIKV nsP2-Chk2 is classified as relatively transient that indicates the reversible nature of the interaction (37,38).

The *in vitro* results showed that AAKi treatment drastically reduced CHIKV infection which was further corroborated in the C57BL/6 mice model. While infected mice displayed moderate disease with progressive weight loss, lethargy, hunched posture, ruffled fur arthritis and immobility, AAKi (2 mg/kg) mitigated these ailments with a lessening in the viral RNA copy number and proteins. The decrease in disease symptoms, nsP2 protein levels and viral RNA copy number, were well supported by the clinical score. Additionally, AAKi was examined against CHIKV infection *ex vivo* in hPBMC-derived monocyte-macrophages. Although CHIKV infection enhanced viral load and E2 positive cells, treatment with AAKi diminished these effects, indicating the importance of these pathways in CHIKV life cycle.

CHIKV infection is capable of controlling cell-cycle arrest in both the G1 and G2 phases, possibly due to Chk1 and Chk2 activation. A number of DNA and RNA viruses also have reported to arrest cell cycle so that they can create a favourable milieu for viral replication. Cell cycle arrest in G1 phase is reported earlier for JEV, Influenza A virus, Severe acute respiratory Syndrome Coronavirus (SARS-CoV), Kaposi’s Sarcoma-Associated Herpesvirus and the murine coronavirus mouse hepatitis virus (MHV), while G2 phase arrest is known for Bocavirus, Epstein-Barr virus and HIV (8,28,39). Nevertheless, there are several unresolved questions in reference to the precise consequences of viral induced cell cycle arrest which needs to be explored further.

As per the findings, it can be summarized that CHIKV infection leads to extensive DNA damage and stimulates both the ATM/Chk2 and ATR/Chk1 signaling pathways. It also appears that both Chk2 and Chk1 play crucial function in CHIKV life cycle. Following CHIKV infection, Chk2 and Chk1 were phosphorylated, which causes cell cycle arrest in both G1 and G2 phases, apoptosis (40) and senescence (41) that further foster efficient viral production (Figure 11).

**Figure 11.**
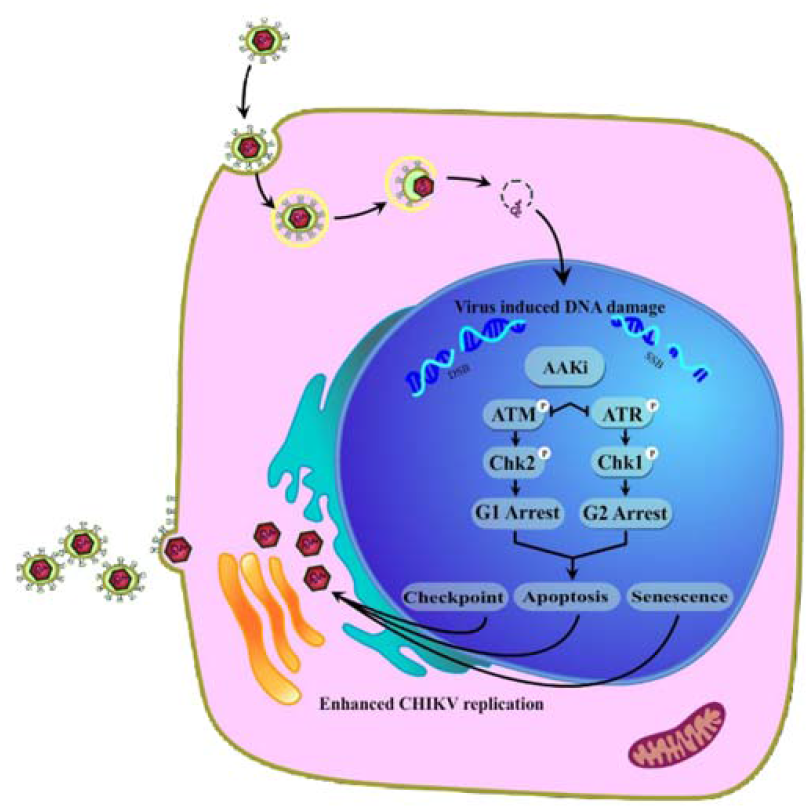
Model portraying modulation of DDR pathways by the CHIKV infection. In the current investigation, it was found that CHIKV infection leads to extensive DNA damage and stimulates both of the ATM/Chk2 and ATR/Chk1 signaling pathways. Inhibition of the crucial host factors linked with DDR pathways causes drastic reduction in the CHIKV infection. Additionally, it also causes cell cycle arrest in both G1 and G2 phases, apoptosis and senescence that further facilitates viral production.

In future, additional investigations are required to decipher the role of amino acids involved in Chk2-nsP2 and Chk1-nsP2 interactions. Creating mutations in these amino acids might help to deepen our understanding about the importance of these interactions in CHIKV infection. The AAKi molecule has shown to be a potential inhibitor of CHIKV infection in mice, however the effect can also be improved further by increasing the dose of the inhibitor to 5mg/kg body weight and or reducing the duration of the drug administration.

In conclusion, this work demonstrated for the first time the mechanistic insight of the induction of DDR pathways by CHIKV and interaction of CHIKV-nsP2 with two crucial host factors, Chk2 and Chk1 for efficient viral infection and this information might contribute to the development of effective therapeutics for the control of the CHIKV infection in future.

## Data availability

The data that support the findings of this study are available from the corresponding author upon reasonable request. Some data may not be made available because of privacy or ethical restrictions.

## SUPPLEMENTAL MATERIAL

SUPPLEMENTARY FILE 1

## Acknowledgments

We are appreciative to Dr. M.M. Parida, DRDE; Gwalior, India for providing CHIKV strain and also thankful to Ms. Shrishti Lama for her help in COMET assay. We are thankful to Mr. Rahul Chatterjee for designing the working model.

